# Beyond hazard identification: Discovering mechanisms of action from a ToxCast chemical screen in zebrafish

**DOI:** 10.64898/2026.09.01.748647

**Authors:** Shayan Shahriar, Daniel A Gorelick

**Affiliations:** Center for Precision Environmental Health, Department of Molecular & Cellular Biology, Baylor College of Medicine, Houston, Texas, USA

## Abstract

Large-scale chemical screens are important tools for hazard identification and chemical prioritization, but they less commonly progress from identifying a phenotype to identifying its mechanism. Here, we used zebrafish embryos to screen 4,657 chemicals from the U.S. EPA ToxCast Phase III library for disruption of embryonic development and advanced selected hits through sequential validation using original library stocks, independently sourced chemicals, concentration-response analysis, transcriptomics, and functional experiments. Of 61 primary hits subjected to repeat testing, 33 reproduced the original phenotype, and four of eight compounds subsequently tested using independently sourced chemicals exhibited reproducible concentration-dependent developmental toxicity. We then investigated purpurin, an understudied anthraquinone pigment that caused pericardial edema, circulation defects, and body-axis abnormalities. Transcriptomic analysis of purpurin-exposed embryos revealed coordinated suppression of pathways involved in calcium regulation, ion transport, and neuronal signaling. Increasing extracellular calcium produced a concentration-dependent rescue of purpurin-induced developmental abnormalities, whereas equivalent magnesium supplementation did not, supporting a role for calcium availability or homeostasis in purpurin developmental toxicity. These results demonstrate that large-scale in vivo toxicity screening can be integrated with independent chemical validation and functional follow-up to move beyond hazard identification toward mechanistic understanding of how environmental chemicals disrupt embryonic development.

## INTRODUCTION

Exposure to environmental chemicals can disrupt embryonic development, leading to defects in growth, organogenesis, and physiological function. Most human birth defects lack a defined cause and are thought to arise from complex combinations of genetic and environmental factors. Although thousands of chemicals are used in industrial, agricultural, and consumer applications, developmental hazard information remains unavailable for many compounds. Testing chemicals individually in animal models is resource-intensive and impractical for the number of compounds requiring evaluation. To address this problem, the U.S. Environmental Protection Agency (EPA) established the Toxicity Forecaster (ToxCast) program to evaluate the biological activity of thousands of chemicals using high-throughput cell-based assays (Judson et al. 2009, 2010). However, in vitro assays cannot reproduce embryonic development in an intact animal and therefore cannot determine whether chemical-induced molecular changes result in developmental abnormalities. Whole-organism models can address this limitation by directly testing chemical effects during embryogenesis.

Zebrafish (*Danio rerio*) embryos provide a vertebrate model in which thousands of chemicals can be tested for effects on development (Pardo-Martin et al. 2010; Peterson et al. 2000). Previous studies used zebrafish to screen chemical libraries and identified compounds that cause embryonic phenotypes (McCollum et al. 2017; Padilla et al. 2012; Raftery et al. 2014; Reif et al. 2016; Truong et al. 2014). Zebrafish toxicology data has also been used to develop computational models that predict developmental toxicity (Balik-Meisner et al. 2018; Green et al. 2021; Thomas et al. 2019; Truong et al. 2016; Zhang et al. 2016, 2017b, 2017a). However, these chemical screens have generally emphasized hazard identification and chemical prioritization, with comparatively limited experimental follow-up to determine how individual chemicals produce developmental phenotypes (Truong et al. 2014).

Here, we screened 4,657 chemicals from the EPA ToxCast Phase III library for effects on zebrafish embryogenesis and subjected hit chemicals to sequential validation using repeat screening, independently sourced compounds, concentration-response analysis, transcriptomics, and functional experiments. Among the reproducible hits, we selected 4-aminoazobenzene and purpurin for further study because the mechanisms underlying their developmental toxicity were unknown. Transcriptomic analysis of 4-aminoazobenzene identified altered neuronal, receptor-mediated, and stress-response pathways and led us to test AHR2 as a candidate mediator. However, AHR2 was not required for developmental toxicity. In contrast, purpurin produced widespread developmental abnormalities and a transcriptional signature implicating calcium-dependent signaling. Extracellular calcium, but not magnesium, rescued purpurin-induced developmental toxicity, supporting a role for altered calcium availability or homeostasis. Together, these experiments establish an approach for advancing hits from toxicology screens from phenotype identification toward mechanisms of developmental toxicity.

## RESULTS

### Chemical screening identifies developmental phenotypes in zebrafish

To identify chemicals that disrupt embryonic development, we screened 4,657 chemicals from the U.S. EPA ToxCast Phase III library using zebrafish embryos (Fig. 1A). We first performed a pilot screen using two 96-well ToxCast plates (plates #0351 and #0338) to select a concentration for the full screen. Three embryos in a single well were exposed to each chemical, and a chemical was classified as a hit when all three embryos exhibited a developmental phenotype. At 2 µM, 1.6% of the tested chemicals met this criterion, providing a screening concentration that yielded a small but detectable proportion phenotypes. We therefore used 2 µM for the full library screen.

**Figure 1.**
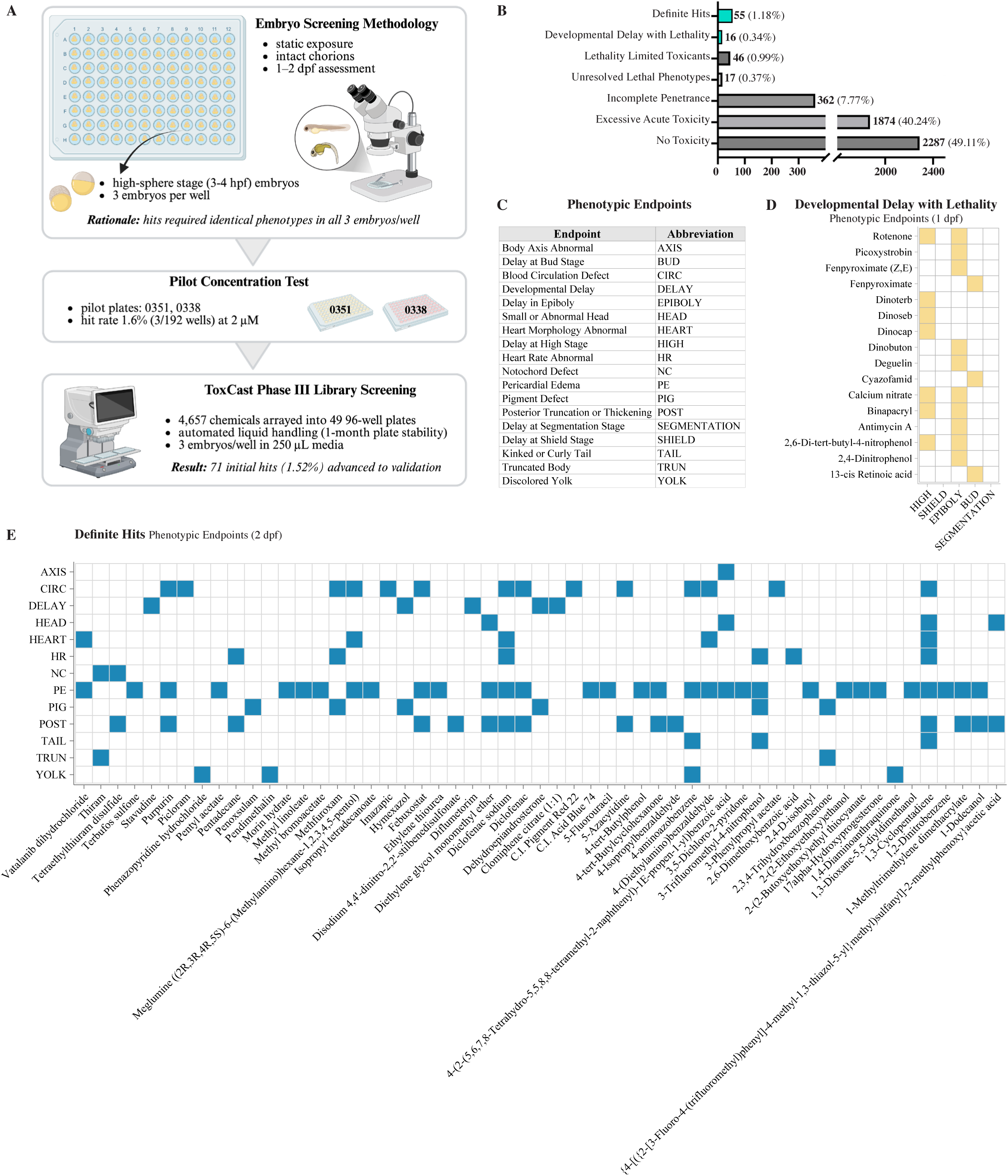
High-throughput chemical toxicity screening in zebrafish. **(A)** Overview of the zebrafish ToxCast chemical screening workflow, including pilot concentration testing and large-scale screening of the Phase III library. **(B)** Classification of 4,657 chemicals screened at 2 µM based on embryo survival and developmental phenotype outcomes. Chemicals were categorized as No Toxicity (all embryos alive and phenotypically normal at 2 dpf), Excessive Acute Toxicity (at least one mortality before 1 dpf screen), Incomplete Penetrance (presence of at least 1 normal embryo at 2 dpf), Unresolved Lethal Phenotypes (all three normal at 1 dpf but dead before 2 dpf assessment), Lethality-Limited Toxicants (all alive at 1 dpf, partial mortality at 2 dpf, alive embryos have phenotypes at 2 dpf), Developmental Delay with Lethality (developmental delay phenotypes evident at 1 dpf, high lethality at 2 dpf), or Definite Hits (all embryos alive and phenotypes exhibited at 2 dpf endpoint). Values indicate the number and percentage of screened chemicals in each category. **(C)** Phenotypic endpoint abbreviations used for developmental toxicity scoring. **(D)** Developmental delay endpoints observed among Early Developmental Hits at 1 dpf. Colored squares indicate developmental arrest at the indicated embryonic stage (x axis) per chemical (y axis). **(E)** Developmental phenotype profiles of Definite Hits at 2 dpf. Colored squares indicate the phenotype endpoint (y axis) for each chemical (x axis).

Embryos were exposed beginning at the sphere stage (3–4 hpf) and evaluated at 1 and 2 days post-fertilization (dpf) for viability and developmental phenotypes (Fig. 1B). We categorized chemicals into 7 groups based on mortality and phenotype penetrance. Chemicals classified as *no toxicity* produced three viable embryos at 2 dpf with no detectable developmental abnormalities and morphology comparable to the DMSO-treated controls. This group represented the largest proportion of the screened library and included 2,287 chemicals (49.11%) (Supplementary Table 2). A second major category, *excessive acute toxicity*, consisted of chemicals for which at least one of the three embryos died before the 1 dpf assessment. Because early lethality reduced the number of embryos available for developmental evaluation, these compounds could not be reliably assessed for later morphological abnormalities and were therefore excluded from subsequent phenotypic classification. This category comprised 1,874 chemicals (40.24%) (Supplementary Table 3). Chemicals classified as *incomplete penetrance* produced three surviving embryos at 1 dpf but retained at least one morphologically normal embryo at 2 dpf. These compounds therefore failed to produce fully penetrant developmental phenotypes across all three embryos and were not advanced as primary screening hits. This category included 362 chemicals (7.77%) (Supplementary Table 4). A small number of compounds were classified as *unresolved lethal phenotypes*, in which all embryos appeared phenotypically normal at 1 dpf but all three embryos died by 2 dpf. Because mortality occurred between the two scoring time points, the developmental abnormalities responsible for lethality could not be resolved morphologically, preventing assignment of a definitive developmental phenotype. 17 chemicals (0.37%) met these criteria (Supplementary Table 5). Another group of compounds was classified as *lethality-limited toxicants*, in which all embryos survived to 1 dpf, one or more embryos died before the 2 dpf assessment, and the remaining surviving embryos displayed clear developmental abnormalities. Although these chemicals induced scorable developmental phenotypes, interpretation was complicated by partial lethality within the well, and they were therefore considered separately from fully penetrant developmental hits. This category contained 46 chemicals (0.99%) (Supplementary Table 6).

Two outcome categories were prioritized for downstream validation. *Developmental delay with lethality* included chemicals that consistently delayed embryonic development at the 1 dpf assessment but resulted in extensive lethality by 2 dpf, limiting evaluation of later morphological phenotypes. 16 chemicals (0.34%) were assigned to this category (Supplementary Table 7). In contrast, *definite hits* consisted of chemicals for which all embryos survived to 2 dpf and displayed developmental abnormalities. 55 chemicals (1.18%) met these criteria and represented the highest-confidence developmental toxicants identified in the primary screen (Supplementary Table 8). Together, the *developmental delay with lethality* and *definite hit* categories yielded 71 primary screening hits. These 71 chemicals produced a total of 18 phenotypic endpoints including developmental delay, body axis abnormalities, blood circulation defects, head defects, heart morphology abnormalities, heart rate abnormalities, notochord defects, pericardial edema, pigment defects, posterior truncation, tail abnormalities, truncated body, and yolk discoloration (Fig. 1C).

Developmental delay phenotypes observed among the *developmental delay with lethality* group were not confined to a single stage of embryogenesis but instead represented arrest or delayed progression across multiple developmental stages, including high, shield, epiboly and bud stages (Fig. 1D). This diversity suggests that chemically induced developmental delay can arise through disruption of distinct developmental processes operating during early embryogenesis rather than through a single common mechanism.

The 55 *definite hits* collectively produced a diverse spectrum of morphological abnormalities at 2 dpf (Fig. 1E). Several phenotypes occurred frequently across multiple chemicals, whereas others were comparatively rare. Pericardial edema occurred frequently, in more than half of the *definite hits*, and was often accompanied by abnormalities in heart morphology, impaired blood circulation, pigment defects, and posterior body malformations. In contrast, phenotypes such as truncated body were observed infrequently. Many compounds produced multiple phenotypic endpoints simultaneously, indicating that individual chemicals frequently produced complex developmental phenotypes rather than a single morphological abnormality.

### Validation of primary screening hits

To determine whether the primary screening hits were reproducible, we re-tested chemicals from the two prioritized hit categories: the 55 *definite hits* and 16 *developmental delay with lethality* compounds, representing 71 candidate developmental toxicants from the primary screen (Fig. 2A). Of these, 61 chemicals (85.9%) were re-validated using the same screening format as the primary assay. The remaining 10 chemicals were identified as hits during subsequent review of the primary screening data after secondary validation experiments had been completed and were therefore not retested.

**Figure 2.**
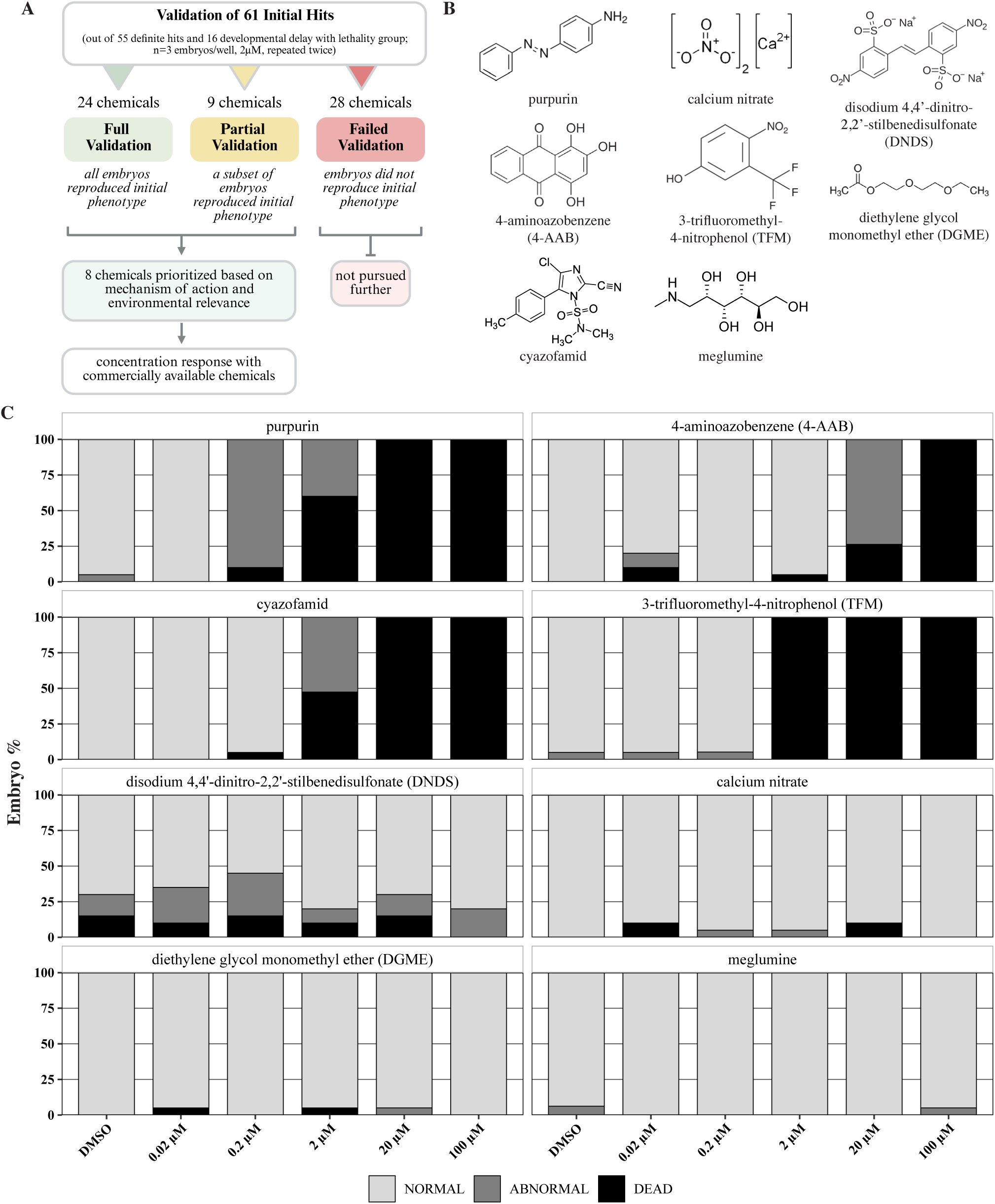
Validation of 71 ToxCast initial hits. **(A)** Validation and prioritization workflow. Sixty-one candidate developmental toxicants identified in the primary screen were retested at 2 µM (n = 3 embryos per well, repeated twice) and classified as full, partial, or failed validation based on phenotype reproducibility. Eight compounds from the fully and partially validated groups were prioritized for further study based on putative mechanism of action, and environmental relevance. **(B)** Chemical structure of 8 prioritized compounds. **(C)** Concentration–response assessment of prioritized compounds using commercially sourced chemicals. Embryos were exposed from 3-4 hpf and evaluated for developmental toxicity at 2 dpf. Each treatment group consisted of 19–21 embryos. Bars indicate the percentage of embryos classified as normal, abnormal, or dead. Abnormal embryos exhibited one or more developmental phenotypic endpoints defined in Figure 1C and Supplementary Figure 1. Purpurin, 4-aminoazobenzene (4-AAB), cyazofamid, and 3-trifluoromethyl-4-nitrophenol (TFM) reproduced developmental toxicity observed in the primary screen, whereas calcium nitrate, DGME, meglumine, and DNDS showed limited or inconsistent phenotypic responses under the conditions tested.

Validation outcomes were classified into three categories. Chemicals were considered *fully validated* if both independent daughter-plate tests reproduced the original phenotype in all exposed embryos, corresponding to six of six embryos affected across the two validation runs. 24 chemicals (39.34%) met this criterion. Chemicals were considered *partially validated* if the original phenotype was reproduced in a subset, but not all, of the six exposed embryos. 9 chemicals (14.75%) met this criterion. Chemicals were classified as *failed validation* if the original phenotype was not reproduced in any exposed embryos or if the phenotype was inconsistent with the primary screen (Supplementary Table 9). The 28 chemicals (45.9%) that failed validation were not pursued further. Together, the full and partial validation groups defined 33 chemicals (54.1%) that reproduced the primary screening phenotype summarized in Table 1.

**Table 1.**
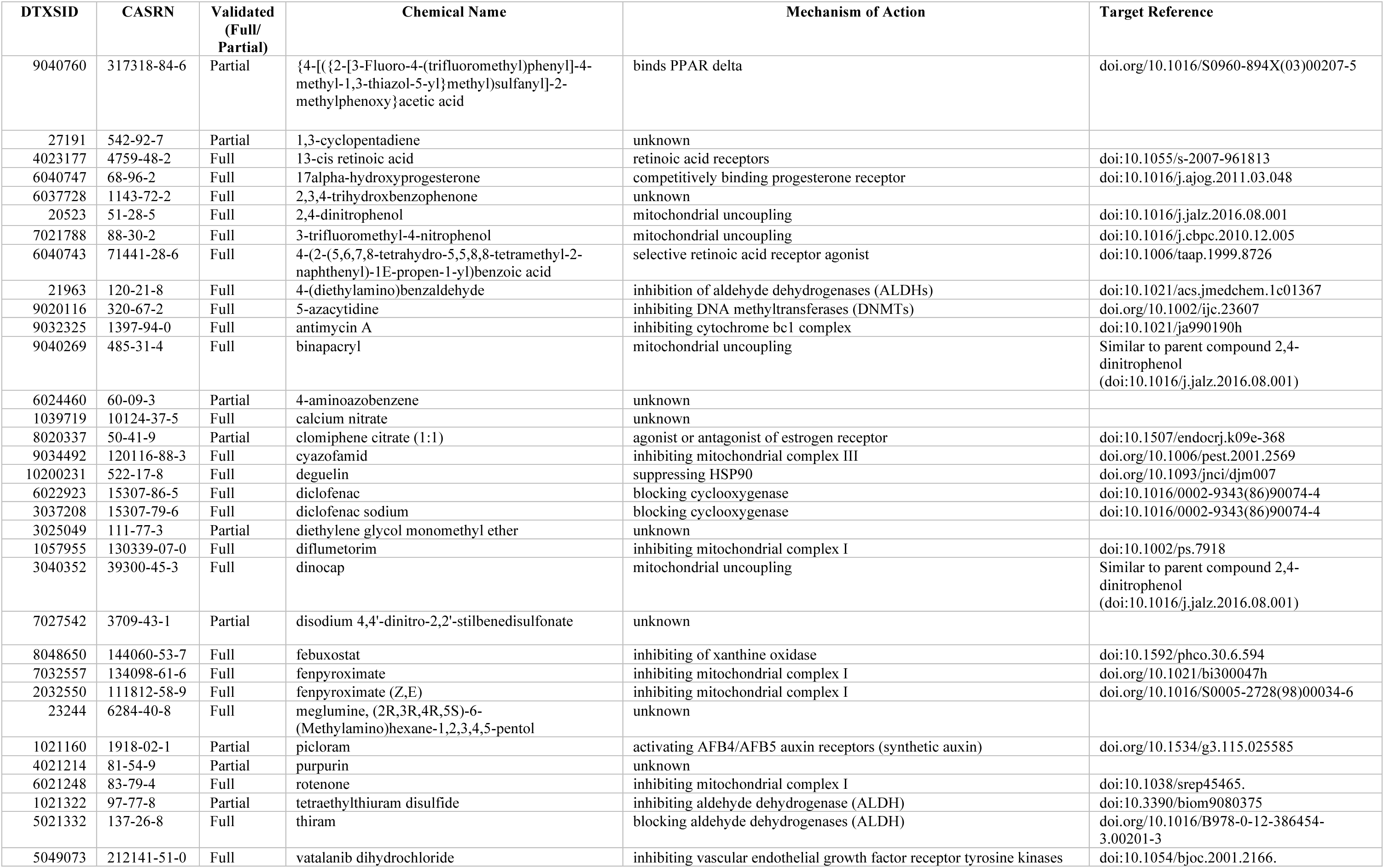
List of 33 validated chemicals identified from 71 initial hits. (comprising 55 definitive hits and 16 compounds associated with developmental delay with lethality)

From the validated chemical set, we selected 8 compounds for concentration-response analysis based on their environmental relevance, commercial availability and lack of knowledge regarding mechanisms of action (Fig. 2B, Table 1). We wanted to determine whether their developmental phenotypes could be reproduced using chemicals obtained independently of the ToxCast library. Priority was given to chemicals with poorly characterized mechanisms underlying developmental toxicity, including calcium nitrate, meglumine, purpurin, 4-aminoazobenzene, diethylene glycol monomethyl ether (DGME), and disodium 4,4′-dinitro-2,2′-stilbenedisulfonate (DNDS). We also selected two environmentally relevant chemicals with established mechanisms of action: 3-trifluoromethyl-4-nitrophenol (TFM), a widely used lampricide (Birceanu et al. 2011) and cyazofamid, a commonly used agricultural fungicide (Mitani et al. 2001). These eight chemicals were obtained from commercial vendors and evaluated independently of the original EPA-sourced ToxCast library. For each chemical and concentration, 19–21 embryos were exposed in 3 mL of treatment medium and scored at 2 dpf as normal, abnormal, or dead (Fig. 2C). Abnormal embryos were defined as those exhibiting one or more developmental phenotypic endpoints described for each chemical (Supplementary Fig. 1, Supplementary Table 10).

Commercial validation separated the eight prioritized chemicals into two groups based on reproducibility. Calcium nitrate, meglumine, diethylene glycol monomethyl ether (DGME), and disodium 4,4′-dinitro-2,2′-stilbenedisulfonate (DNDS) failed to reproducibly induce developmental toxicity across the tested concentration range compared with DMSO controls and were therefore excluded from further investigation. In contrast, purpurin, 4-aminoazobenzene, cyazofamid, and 3-trifluoromethyl-4-nitrophenol (TFM) exhibited concentration-dependent toxicity.

To more precisely define the developmental toxicity profiles of the four reproducible chemicals, purpurin, 4-aminoazobenzene, cyazofamid, and 3-trifluoromethyl-4-nitrophenol (TFM), we performed concentration-response experiments in six-well plates (Fig. 3; Supplementary Table 11). Unlike the preliminary commercial validation, this experiment consisted of three independent biological replicates derived from separate embryo clutches. For each concentration, 20 embryos were exposed between the sphere and 50% epiboly stages and scored at 1 and 2 days post fertilization (dpf) for developmental abnormalities and mortality.

**Figure 3.**
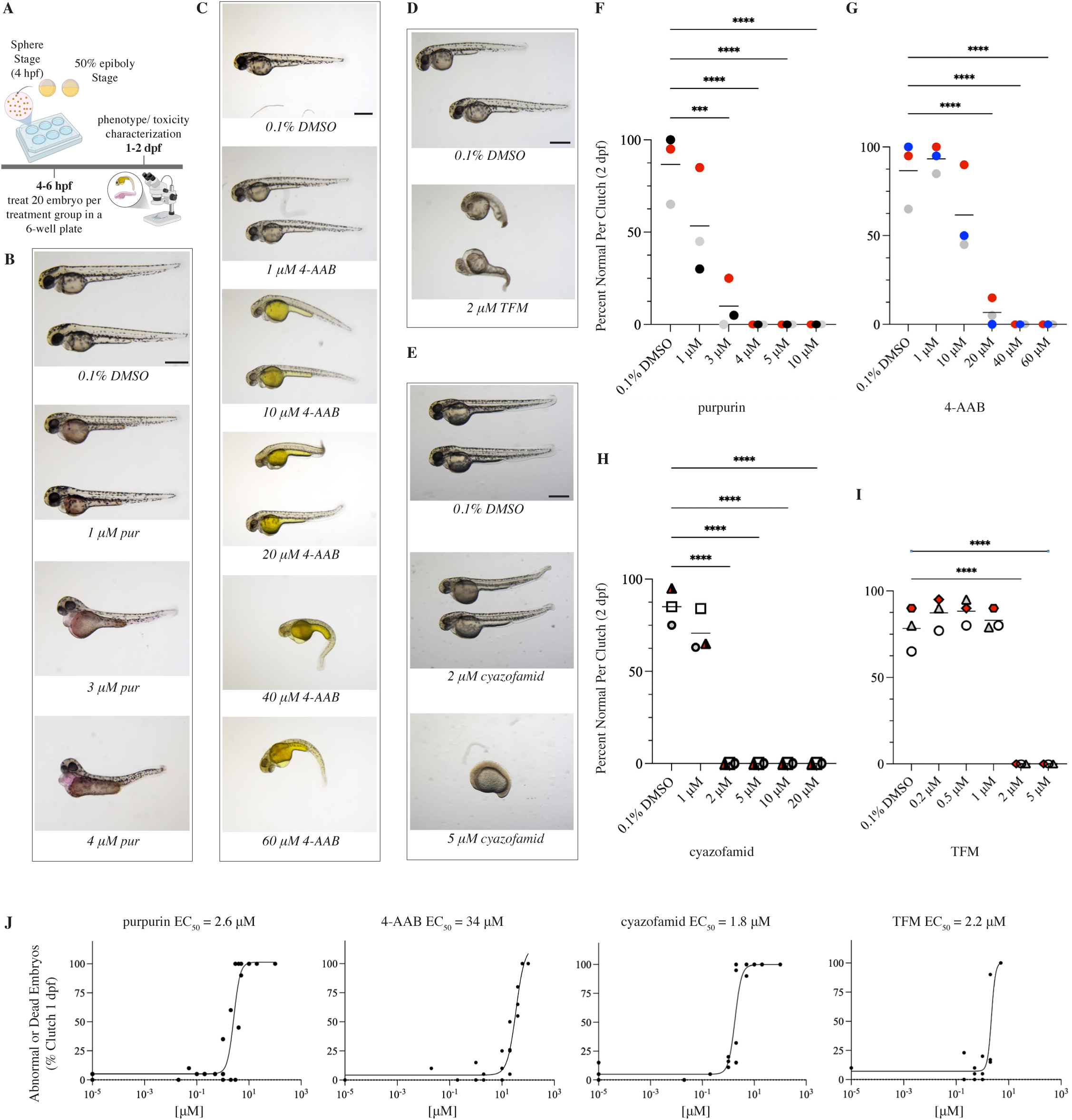
Developmental toxicity of prioritized ToxCast hits in zebrafish embryos. **(A)** Experimental workflow for zebrafish embryo exposure and phenotypic assessment. **(B–E)** Representative images of embryos exposed to increasing concentrations of purpurin, 4-aminoazobenzene (4-AAB), 3-trifluoromethyl-4-nitrophenol (TFM), and cyazofamid from 4–6 hours post-fertilization (hpf) and imaged at 2 days post-fertilization (dpf). Scale bar, 0.5 mm. **(F–I)** Quantification of developmental toxicity as percent normal embryos per clutch at 2 dpf. Each point represents one clutch of 20–21 embryos (n = 20–21), with three biological replicates (N = 3) indicated by different symbols or colors. Horizontal lines denote mean values. Statistical significance was determined by one-way ANOVA followed by Dunnett’s multiple comparisons test (*** adjusted P < 0.001, **** adjusted P < 0.0001; ns). **(J)** Concentration–response curves for developmental toxicity induced by each compound. Each point represents one clutch scored for abnormal or dead phenotypes at 1 dpf (N = 3 biological replicates per concentration). Nonlinear regression was used to estimate EC50 values.

Purpurin consistently induced developmental abnormalities characterized primarily by pericardial edema, body axis defects, posterior truncation, yolk discoloration, and impaired circulation (Fig. 3B). The frequency of abnormal and dead embryos increased sharply between 1 and 3 µM, with extensive mortality observed at higher concentrations (Fig 3F, Supplementary Table 11). Concentration-response data were used to calculate the half-maximal effective concentration (EC₅₀) for purpurin to be 2.6 µM (Fig. 3J; Supplementary Table 12).

4-Aminoazobenzene produced a distinct developmental phenotype dominated by tail abnormalities, body axis defects, pericardial edema, and yolk discoloration (Fig. 3C). Yolk discoloration was also observed following purpurin exposure, consistent with the dye properties of both compounds. Approximately 60% of embryos remained morphologically normal at 10 µM, whereas developmental abnormalities increased markedly at 20 µM and became nearly fully penetrant at 40 µM, where mortality was also frequently observed (Fig. 3G; Supplementary Table 11). The EC₅₀ for 4-aminoazobenzene was 34 µM (Fig. 3J; Supplementary Table 12).

Cyazofamid and TFM exhibited comparatively steep concentration-response relationships. Approximately 70% of embryos exposed to 1 µM cyazofamid remained phenotypically normal, whereas exposure to 2 µM resulted in nearly all embryos becoming abnormal or dead. The predominant phenotypes included developmental delay, body axis defects, and tail deformities (Fig. 3E, H; Supplementary Table 11). TFM produced a similar concentration-response profile. A greater proportion of embryos (approximately 83%) exhibited normal development at 1 µM before progressing to fully penetrant developmental toxicity or lethality at higher concentrations. Characteristic TFM-induced phenotypes included body axis defects, tail abnormalities, pericardial edema, and heart defects (Fig. 3D, I). The EC₅₀ values for cyazofamid and TFM were 1.8 µM and 2.2 µM, respectively (Fig. 3J; Supplementary Table 12).

Validation using commercially supplied compounds reduced the eight prioritized screening hits to four chemicals with reproducible concentration-dependent developmental toxicity: purpurin, 4-aminoazobenzene, cyazofamid, and TFM. Because the mechanisms of action of cyazofamid and TFM are known, with cyazofamid inhibiting mitochondrial complex III (Mitani et al. 2001) and TFM acting as a mitochondrial uncoupler (Birceanu et al. 2011), we focused subsequent experiments on purpurin and 4-aminoazobenzene.

### 4-Aminoazobenzene disrupts neuronal and receptor-mediated signaling during development

To investigate the molecular basis of 4-aminoazobenzene-induced developmental toxicity, zebrafish embryos were exposed to 40 µM 4-aminoazobenzene beginning at 50% epiboly. Whole-embryo RNA was isolated at 2 dpf from a pooled sample of 20 phenotypically affected, live embryos for transcriptomic analysis (Fig. 4A). Differential expression analysis identified 3,259 significantly altered genes (FDR < 0.05), including 1,485 upregulated and 1,774 downregulated genes (Fig. 4B), indicating a broad transcriptional response to 4-aminoazobenzene exposure (Supplementary Table 15).

**Figure 4.**
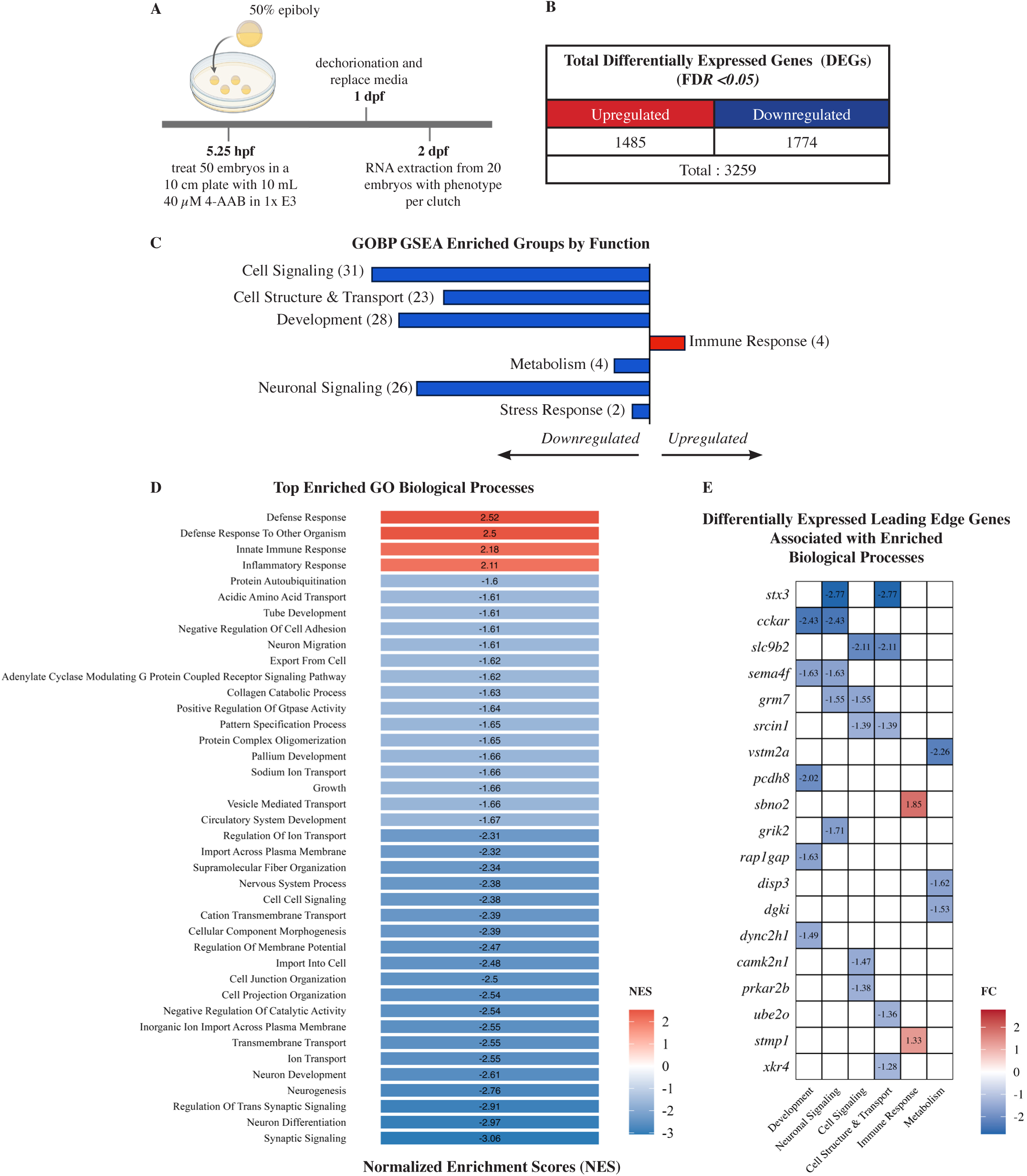
Transcriptomic response to 4-aminoazobenzene (4-AAB) exposure in zebrafish embryos. **(A)** Experimental workflow for RNA-seq analysis. Embryos were treated at 50% epiboly (5.25 hpf) with 40μM 4-AAB, and maintained until RNA extraction at 2 days post-fertilization (2 dpf). Each sample consisted of 20 embryos per clutch across three biological replicates (N=3, n=20). **(B)** Summary of differentially expressed genes (DEGs) identified by RNA-seq following 4-AAB exposure (FDR < 0.05), showing the number of upregulated (1485) and downregulated (1774) genes (total = 3259) compared to 0.1% DMSO-treated controls. **(C)** Functional categorization of significantly enriched (FDR < 0.25) Gene Ontology Biological Process (GOBP) terms based on gene set enrichment analysis (GSEA). Enriched pathways were grouped into biological categories. Numbers in parentheses indicate the number of GOBP terms per category. Bar direction indicates the dominant direction of enrichment, with negative normalized enrichment scores (NES) representing downregulation and positive NES representing upregulation. **(D)** Top enriched GOBP terms ranked by normalized enrichment score (NES). Red indicates positively enriched (upregulated) pathways, and blue indicates negatively enriched (downregulated) pathways. **(E)** Heatmap of selected differentially expressed leading edge genes associated with enriched biological processes. Color represents fold change (FC), with blue indicating downregulation and red indicating upregulation.

To determine the biological processes most affected by 4-aminoazobenzene exposure, gene set enrichment analysis (GSEA) was performed using Gene Ontology Biological Process (GOBP) annotations, which were subsequently grouped into seven broad functional categories (Supplementary Table 13). Although both positively and negatively enriched biological processes were identified, downregulated pathways predominated and were primarily associated with neuronal signaling, development, cell signaling, and cell structure and transport (Fig. 4C). In contrast, the relatively few positively enriched biological processes were largely restricted to immune response.

The only positively enriched biological processes were associated with defense and innate immune responses, whereas the vast majority of significantly enriched biological processes were negatively enriched (Fig 4D, Supplementary Table 16). These downregulated processes encompassed multiple aspects of vertebrate development, with the five most significantly repressed biological processes converging on neuronal development and function, including neurogenesis, neuron differentiation, and synaptic signaling.

To identify genes contributing to these enriched processes, we examined the top five leading-edge genes within each functional category (Fig 4E, Supplementary Table 14). Four genes represented across multiple functional categories—*cckar, rap1gapa, prkar2b,* and *grm7**—***share a common role in G protein-coupled receptor (GPCR)-mediated signaling. GPCRs constitute the largest and most diverse family of membrane receptors and regulate numerous biological processes, including embryonic development, neuronal signaling, cell differentiation, calcium and ion homeostasis, and tissue morphogenesis (Rosenbaum et al. 2009). The coordinated downregulation of multiple GPCR-associated genes suggests receptor-mediated signaling as a process disrupted by 4-aminoazobenzene.

To further refine the broad biological processes identified by GOBP analysis, we performed GSEA using KEGG and Hallmark pathway databases (Fig. 5A). The most significantly downregulated canonical pathway was *neuroactive ligand-receptor interaction*, consistent with neuronal and receptor-mediated signaling. Additional downregulated pathways included *calcium signaling* and *long-term depression*, both of which are associated with synaptic function and neuronal signaling. *Vascular smooth muscle* contraction was also significantly downregulated, providing a potential link to the cardiovascular phenotypes observed following 4-aminoazobenzene exposure. In contrast, the most significantly upregulated pathways included interferon-γ response, hypoxia, and the unfolded protein response, indicating activation of cellular stress and immune-related signaling pathways.

**Figure 5.**
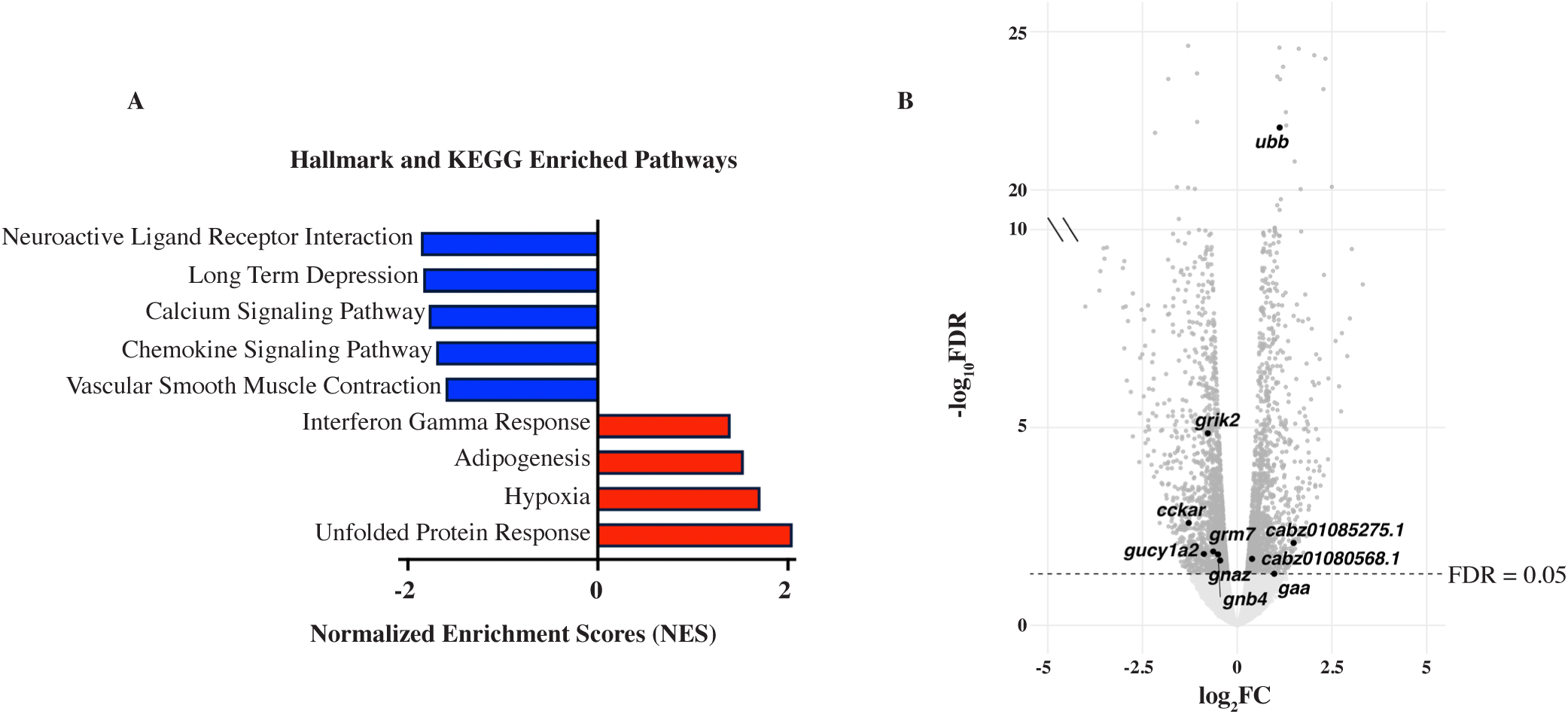
4-aminoazobenzene–induced pathway enrichment. **(A)** Significantly enriched (FDR < 0.25) pathways from KEGG and Hallmark gene sets identified by gene set enrichment analysis (GSEA). Pathways are ranked by normalized enrichment score (NES). Negative NES values indicate downregulation, and positive NES values indicate upregulation, highlighting coordinated repression of neuronal and calcium signaling pathways alongside modulation of stress and immune-related pathways. **(B)** Volcano plot showing differential gene expression following 4-AAB exposure. Each point represents a gene plotted by log_2_ fold change (FC) and −log_10_(FDR). Dotted line indicates significance thresholds (FDR = 0.05). Highlighted genes represent leading-edge genes from significantly enriched KEGG and Hallmark pathways that meet significance (FDR < 0.05) threshold.

Examination of the leading-edge genes driving these enriched pathways revealed a recurring pattern of receptor-mediated signaling disruption (Fig. 5B). Several significantly downregulated genes, including *cckar*, *grm7*, *gnas*, and *gnb4*, are involved in GPCR-mediated signaling, supporting the coordinated repression of the neuroactive ligand-receptor interaction pathway. Likewise, downregulation of genes associated with calcium signaling further reinforced the suppression of neuronal communication identified across both GOBP and KEGG analyses. In contrast, the upregulated genes reflected activation of stress responses, with *ubb* consistent with enhanced protein quality-control pathways and *gaa* suggesting altered glycogen metabolism accompanying cellular stress. Together, the pathway- and gene-level analyses led us to hypothesize that 4-aminoazobenzene disrupts development by suppressing receptor-mediated signaling while activating cellular stress-response pathways.

We next examined individual differentially expressed genes for candidate receptors and pathways that could mediate 4-aminoazobenzene toxicity. Among the most strongly induced genes was *cyp1a*, a canonical transcriptional target of AHR signaling. *cyp1a* expression increased 12.4-fold following 4-aminoazobenzene exposure (Supplementary Fig. 2A,B). This strong induction prompted us to test whether zebrafish AHR2, a functional ortholog of human AHR, contributes to the developmental toxicity of 4-aminoazobenzene. We exposed maternal-zygotic *ahr2* mutant (MZ*uab147*) and wild-type embryos to 40 µM 4-aminoazobenzene and assessed developmental toxicity at 2 dpf. *ahr2* mutations did not reduce the frequency of abnormal or dead embryos following 4-aminoazobenzene exposure (Supplementary Fig. 2C, Supplementary Table 20). Thus, despite induction of the AHR-responsive gene *cyp1a*, AHR2 was not required for the overt developmental toxicity caused by 4-aminoazobenzene under these exposure conditions.

### Purpurin disrupts calcium signaling

To investigate the molecular mechanisms of purpurin-induced developmental toxicity, zebrafish embryos were exposed to 3 µM purpurin beginning at 50% epiboly. RNA was isolated at 2 dpf from pooled samples of 20 phenotypically affected, live embryos for transcriptomic analysis (Fig. 6A). Differential expression analysis identified 4,072 significantly altered genes (FDR < 0.05), including 1,854 upregulated and 2,218 downregulated genes (Fig. 6B), indicating a transcriptional response dominated by gene repression (Supplementary Table 17).

**Figure 6.**
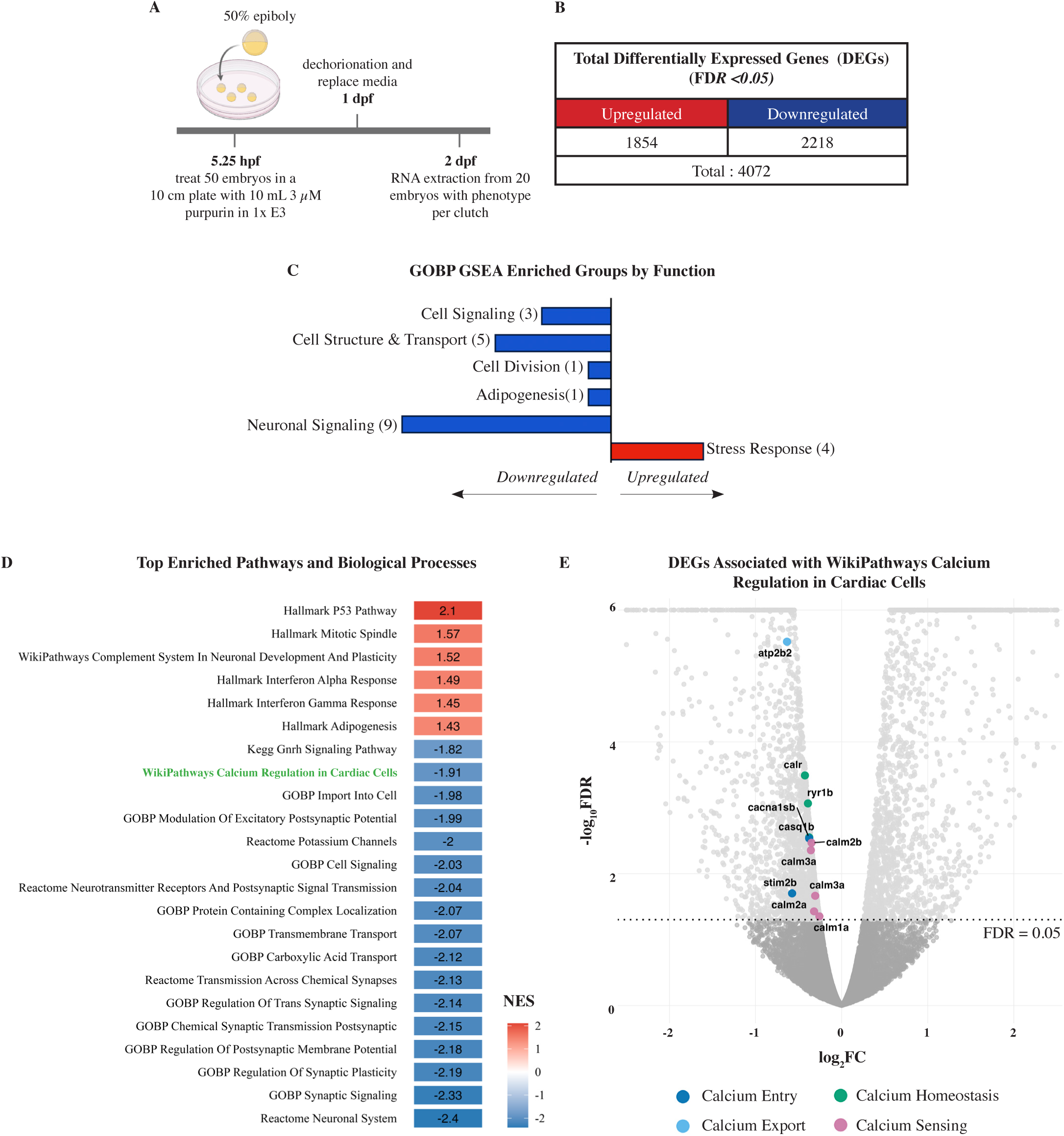
Transcriptomic response to purpurin exposure in zebrafish embryos. **(A)** Experimental workflow for RNAseq. Embryos were treated at 50% epiboly (5.25 hpf) with 3 μM purpurin, and maintained until RNA extraction at 2 days post fertilization (2 dpf). Each sample consisted of 20 embryos per clutch (n=20) across three biological replicates (N=3). **(B)** Summary of differentially expressed genes (DEGs) identified by RNA-seq following purpurin exposure (FDR < 0.05), showing the number of upregulated (1854) and downregulated (2218) genes (total = 4072) compared to 0.1% DMSO-treated controls. **(C)** Functional categorization of significantly enriched pathways identified by gene set enrichment analysis (GSEA) across multiple databases (GOBP, KEGG, Reactome, Hallmark, and WikiPathways). Enriched pathways were grouped into major biological categories. Numbers in parentheses indicate the number of pathways per category. Bar direction indicates the dominant direction of enrichment, with negative normalized enrichment scores (NES) representing downregulation and positive NES representing upregulation. **(D)** Top enriched pathways and biological processes ranked by normalized enrichment score (NES). Red indicates positively enriched (upregulated) pathways, and blue indicates negatively enriched (downregulated) pathways. The *WikiPathways Calcium Regulation in Cardiac Cells* pathway is highlighted in green. **(E)** Volcano plot showing differential gene expression following purpurin exposure. Each point represents a gene plotted by log_2_ fold change (FC) and −log_10_(FDR). Differentially expressed genes associated with the WikiPathways Calcium Regulation in Cardiac Cells pathway are highlighted and color-coded by functional group (calcium entry, calcium export, calcium homeostasis, and calcium sensing). The horizontal dotted line indicates the significance threshold (FDR = 0.05).

To identify the biological pathways most affected by purpurin exposure, gene set enrichment analysis (GSEA) was performed using Gene Ontology Biological Process (GOBP), KEGG, Reactome, Hallmark, and WikiPathways databases. Significantly enriched pathways were manually grouped into six major functional categories (Supplementary Table 18). Similar to 4-aminoazobenzene, negatively enriched pathways predominated and were primarily associated with neuronal signaling and cell structure & transport, whereas positively enriched pathways were largely restricted to stress-response processes (Fig. 6C).

Examination of the highest-ranking enriched pathways further highlighted a diverse transcriptional response (Fig. 6D). The most significantly upregulated pathways included interferon-α response, interferon-γ response, and p53 signaling, consistent with activation of cellular stress-response programs. In contrast, the most significantly downregulated pathways converged on neuronal communication and ion transport, including neuronal system, synaptic signaling, regulation of synaptic plasticity, neurotransmitter receptor and postsynaptic signaling, potassium channel activity, and cell signaling. Among these, the WikiPathways *Calcium Regulation in Cardiac Cells* pathway was of particular interest because purpurin consistently induced severe pericardial edema and circulation defects. Because of the known calcium-chelating activity of purpurin and other anthraquinone dyes (Lee et al. 2016; Macallum 1925), these findings led us to hypothesize that disruption of calcium homeostasis contributes to purpurin-induced developmental toxicity.

To further investigate the WikiPathways *Calcium Regulation in Cardiac Cells* pathway, we examined the leading-edge genes contributing to its negative enrichment, particularly with direct calcium-related functions (Fig. 6E). These genes were uniformly downregulated and represented several complementary aspects of calcium regulation, including calcium entry (*cacna1sb*, *stim2b*), calcium export (*atp2b2*), calcium homeostasis (*calr, ryr1b*), and calcium sensing (*calm1a*, *calm2a*, *calm2b*, *calm3a*). The prominence of downregulated calmodulin genes was consistent with effects on not only calcium transport but also calcium-dependent signal transduction. Consistent with this pattern, GSEA showed that the *Calcium Regulation in Cardiac Cells* pathway was enriched toward the downregulated end of the ranked transcriptome (Supplementary Fig. 3A). We next examined additional leading-edge genes from this pathway that were not classified as direct calcium-regulatory genes (Supplementary Fig. 3B). Most of these genes were also downregulated and included several GPCR-associated signaling components, such as *gng13b*, *adcy7*, *rgs20*, *rgs17*, *gnb5b*, and *gng3*, as well as multiple sodium/potassium transport genes. A small number of upregulated genes were also present, but these were primarily associated with intracellular protein localization rather than direct calcium signaling. Together, these results led us to hypothesize that purpurin disrupts calcium-dependent signaling and ion transport during development.

Because this transcriptomic signature implicated calcium dysregulation as a potential mechanism of purpurin toxicity, we tested whether increasing extracellular calcium in growth media could rescue the developmental phenotype. Embryos were exposed to purpurin in standard E3 medium supplemented with increasing concentrations of calcium, while magnesium supplementation served as a control divalent cation (Fig. 7A). Purpurin treatment in standard E3 medium produced severe developmental abnormalities, whereas increasing extracellular calcium resulted in a concentration-dependent rescue of embryo morphology (Fig. 7B). Quantification across biological replicates demonstrated a corresponding reduction in developmental toxicity, with the highest calcium concentration reducing abnormal embryos to near-control levels (Fig. 7C, Supplementary Table 19). In contrast, equivalent magnesium supplementation failed to improve morphology or reduce toxicity at any concentration tested. Together, these findings show that increasing extracellular calcium, but not magnesium, reduces purpurin-induced developmental toxicity and support a role for calcium availability or homeostasis in the developmental response to purpurin.

**Figure 7.**
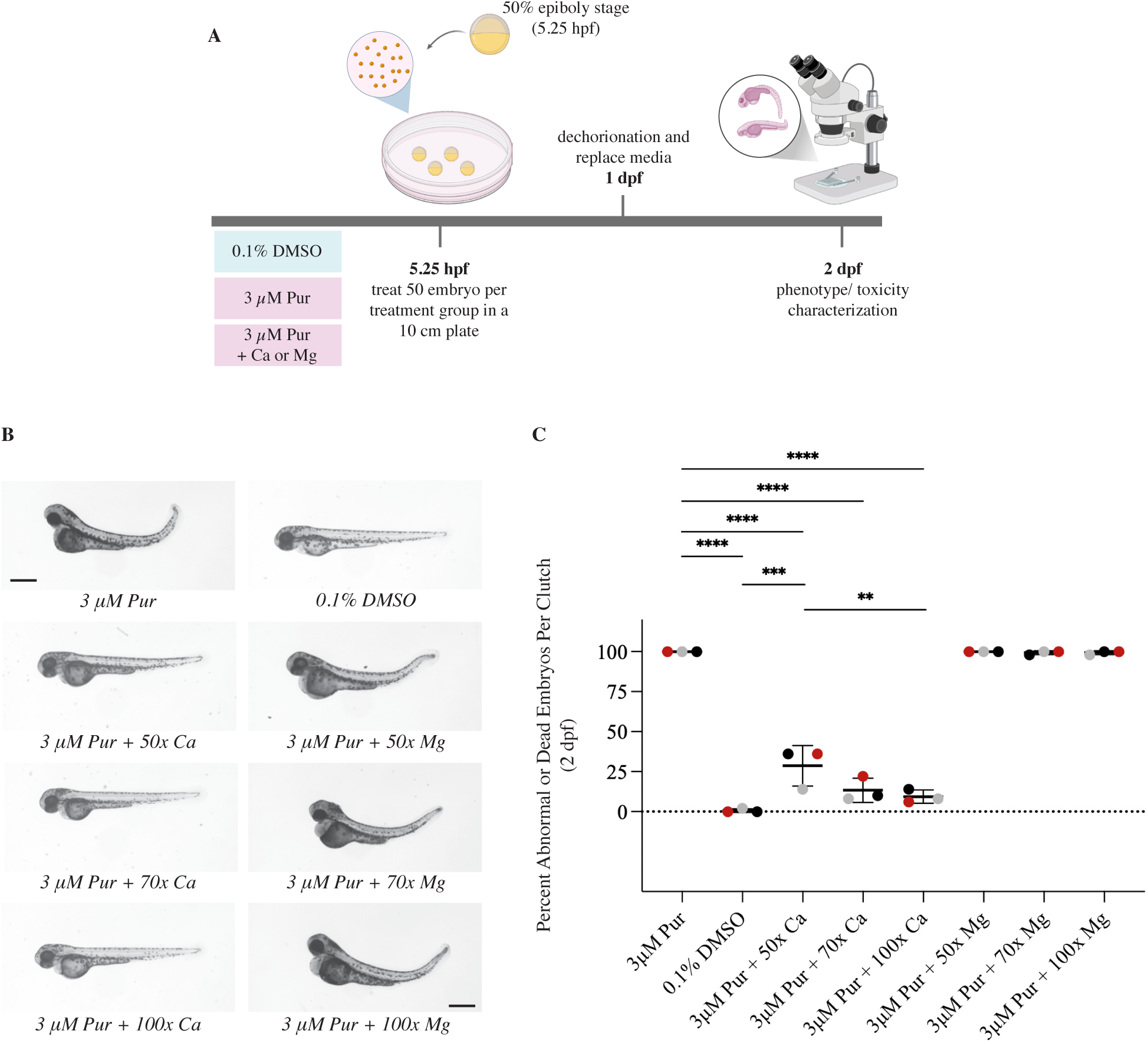
Calcium, but not magnesium, enriched growth media rescues purpurin-induced developmental toxicity in zebrafish embryos. **(A)** Experimental workflow for calcium and magnesium rescue experiments. Embryos were treated at 50% epiboly (5.25 hpf) with 3 μM purpurin in standard E3 medium (containing 0.33 mM Ca²^+^/Mg²^+^) or in E3 supplemented with increasing concentrations of calcium or magnesium. Treatments therefore represent elevated extracellular conditions relative to baseline media. Magnesium was included as a control divalent cation. Each treatment group consisted of 50 embryos per clutch (n=50), with three independent biological replicates (N = 3). **(B)** Representative images of embryos at 2 days post fertilization (2 dpf) following treatment. Purpurin exposure in standard E3 medium induces abnormal morphology, while increasing calcium results in a concentration-dependent rescue of the phenotype. In contrast, magnesium supplementation does not rescue the phenotype. Scale bar, 0.5mm. **(C)** Quantification of developmental toxicity following increased calcium and magnesium in growth media. Data are presented as the percentage of abnormal or dead embryos per clutch at 2 dpf. Each point represents one biological replicate (distinct colors), with mean ± SD indicated. Elevated calcium significantly reduces purpurin-induced toxicity in a concentration-dependent manner, whereas magnesium does not rescue. Statistical significance was determined by one-way ANOVA with Dunnett’s multiple comparisons test (**** adjusted P < 0.0001, *** adjusted P < 0.001, ** adjusted P < 0.01).

## DISCUSSION

The U.S. EPA ToxCast program has generated an extensive resource for prioritizing environmental chemicals based on in vitro bioactivity (Dix et al. 2007; Judson et al. 2010), and previous zebrafish screens have extended this approach by identifying ToxCast chemicals that disrupt embryonic development (McCollum et al. 2017; Padilla et al. 2012; Raftery et al. 2014; Reif et al. 2016; Truong et al. 2014). However, identifying a chemical that produces a developmental phenotype is only the first step toward understanding its toxicity. We identified 71 primary hits, but sequential re-testing with original library stocks and independently sourced chemicals substantially reduced the number of reproducible toxicants. Of eight chemicals subsequently tested using independently sourced compounds, only four—purpurin, 4-aminoazobenzene, cyazofamid, and TFM—reproduced concentration-dependent developmental toxicity. Because their primary molecular actions of cyazofamid and TFM are already well established (Birceanu et al. 2011; Huerta et al. 2020; Mitani et al. 2001; Niblett and Ballantyne 1976), we prioritized purpurin and 4-aminoazobenzene for further investigation. These experiments produced different outcomes: purpurin generated a testable hypothesis involving calcium signaling that was supported by selective rescue with extracellular calcium, whereas 4-aminoazobenzene produced broad transcriptional changes but its initiating mechanism remained unresolved. Together, these findings illustrate the distinction between identifying a developmental hazard and experimentally determining how a chemical disrupts development.

Among the validated developmental toxicants, purpurin is a naturally occurring anthraquinone pigment isolated from *Rubia* plant species and historically used as a dye. It is also used as a fluorescent marker of mineralized tissues, including in zebrafish (Yao et al. 2017). Despite these applications, its effects during early vertebrate development are poorly characterized. One study demonstrated that purpurin inhibits adipocyte-derived leucine aminopeptidase (ERAP1) in vitro and disrupts angiogenesis in zebrafish embryos, but did not examine broader developmental phenotypes or their underlying mechanisms (Park et al. 2014). Purpurin has been detected in madder-dyed textiles, and anthraquinones from madder have been considered potential sources of occupational or consumer exposure (Clementi et al. 2007; Jäger et al. 2006), although purpurin-specific human exposure levels have not been established. Purpurin can also be absorbed following dermal or oral exposure in experimental animals (Gao et al. 2016; Lin et al. 2025). Because the concentrations used in our zebrafish experiments represent nominal water concentrations rather than internal embryo concentrations, they cannot be directly compared with exposure measurements in mammals. Our findings therefore establish developmental toxicity of purpurin at low-micromolar external concentrations but do not establish the relevance of these concentrations to human exposure.

Purpurin exposure caused widespread developmental abnormalities accompanied by altered expression of genes and pathways involved in ion transport and calcium regulation. The coordinated downregulation of genes involved in calcium entry, export, homeostasis, and sensing, together with previous evidence that purpurin interacts with calcium, led us to hypothesize that altered calcium availability or homeostasis contributes to purpurin-induced developmental toxicity. Increasing extracellular calcium produced a concentration-dependent rescue of the developmental phenotype, whereas equivalent magnesium supplementation did not, supporting a specific role for calcium. However, these experiments do not distinguish whether extracellular calcium restores calcium availability or signaling within the embryo, directly interacts with purpurin in the exposure medium and reduces its bioavailability, or acts through both mechanisms. Measurements of intracellular calcium and direct assessment of purpurin– calcium interactions under the exposure conditions will be needed to distinguish these possibilities.

We also investigated 4-aminoazobenzene (4-AAB), an aromatic azo compound used in dye and pigment production and studied for its metabolism, mutagenicity and chemical carcinogenesis (Delclos et al. 1984; Kojima et al. 1992; National Center for Biotechnology Information (2026); Sasaki et al. 1997), but whose effects during vertebrate embryonic development remain poorly characterized. In zebrafish, 4-AAB produced concentration-dependent developmental toxicity with an EC₅₀ of 34 µM. Transcriptomic analysis identified suppression of neuronal, GPCR-mediated, and calcium-signaling pathways, together with activation of immune and cellular stress responses. These changes generated several candidate mechanisms but did not identify an initiating event that could account for the developmental phenotype.

One of the strongest transcriptional responses was induction of *cyp1a*, a canonical AHR target, prompting us to test whether AHR2 was required for 4-AAB toxicity. Mutation of *ahr2* did not reduce the developmental abnormalities or lethality following 4-AAB exposure, demonstrating that AHR2 was not required for the overt developmental toxicity observed under these conditions. Thus, despite a strong AHR-associated transcriptional response, the initiating mechanism of 4-AAB remains unresolved. This result illustrates an important limitation of using transcriptomic responses to infer mechanism: transcriptomics can generate hypotheses, but functional experiments are required to determine whether an implicated pathway is necessary for toxicity.

Because our transcriptomic analyses were performed using whole embryos collected at a single developmental time point, the observed changes may include both primary responses to chemical exposure and secondary consequences of developmental abnormalities. Earlier time points and tissue- or cell-specific analyses could help distinguish initiating molecular events from downstream responses.

The sequential validation process also highlighted the importance of independently confirming screening hits. Several primary hits failed to reproduce during repeat testing, and four of the eight prioritized chemicals failed to reproduce developmental toxicity when independently sourced compounds were tested. Differences between library and independently sourced chemicals could arise from chemical identity, purity, stability, storage, or other factors that cannot be distinguished from the present experiments. Our results emphasize the importance of confirming screening hits before pursuing their mechanisms.

The use of a single 2 µM concentration also limited the phenotypes that could be identified. At this concentration, 1,874 chemicals (40.24%) caused at least one embryo to die before 1 dpf and therefore could not be evaluated for later developmental abnormalities. Future studies could test these chemicals at lower concentrations to determine whether specific developmental phenotypes emerge when acute lethality is reduced. Additionally, screening for gross morphological phenotypes does not detect molecular or physiological effects that occur without visible developmental abnormalities, such as subtle changes in the vasculature. Embryos also remained within the chorion during screening. Because effects of the chorion on chemical uptake are compound-dependent and internal concentrations were not measured, the nominal 2 µM exposure does not represent equivalent embryonic exposure across chemicals. These limitations mean that the chemicals identified here likely represent only a subset of the developmental toxicants present within the library rather than an exhaustive inventory.

In summary, high-throughput phenotypic screening can identify chemicals that disrupt embryonic development, but a screening hit alone does not establish either reproducibility or mechanism. Moving from hazard identification toward mechanism requires independent confirmation of screening phenotypes, generation of specific molecular hypotheses, and experimental testing of those hypotheses. Our results illustrate both outcomes of this process: a mechanistic hypothesis supported experimentally for purpurin, and an unresolved initiating mechanism for 4-AAB.

## MATERIALS AND METHODS

### Zebrafish husbandry and embryo collection

Wild-type AB zebrafish were maintained at 28.5°C under a 14 h light/10 h dark photoperiod in a recirculating aquatic system (Tecniplast USA) at the Baylor College of Medicine Zebrafish Research Facility (Westerfield, M. 2020). Adult male and female zebrafish were placed in static breeding tanks separated by a divider the day before spawning. The divider was removed the following morning to allow natural mating, and embryos were collected within 30 min of spawning. Embryos were maintained in 1x E3 media (prepared by diluting a 60× stock containing 17.2 g NaCl, 0.76 g KCl, 2.9 g CaCl₂·2H₂O, and 2.39 g MgSO₄ in 1 L Milli-Q water to a final volume of 60 L with Milli-Q water) and incubated at 28.5°C. Depending on the experimental design, embryos were transferred to 96-well plates, 6-well plates, or 10-cm Petri dishes for chemical exposure and subsequent analyses. All procedures were approved by the BCM Institutional Animal Care and Use Committee.

### Chemical library and reagents

The U.S. Environmental Protection Agency (EPA) ToxCast Phase III chemical library was obtained from the EPA for primary high-throughput screening. The library consisted of 49 96-well source plates containing 4,657 unique compounds dissolved in dimethyl sulfoxide (DMSO) at a stock concentration of 20 mM. Forty-eight plates each contained 96 compounds (one compound per well), while the final plate contained the remaining 49 compounds.

Assay-ready daughter plates were generated at the Institute of Biosciences and Technology (IBT), Texas A&M University, Houston, Texas, using automated liquid handling. Nanoliter volumes of each compound were dispensed directly from the 20 mM source plates into 96-well assay plates, heat-sealed, and stored at −80°C until use. The dispensing volume was calculated to achieve a final chemical concentration of 2 μM after the addition of embryos in 250 μL E3 medium.

Commercially sourced chemicals used for concentration-response validation and mechanistic studies were purchased from MilliporeSigma unless otherwise indicated. Calcium chloride (CaCl₂) and magnesium sulfate (MgSO₄) used for rescue experiments were also obtained from MilliporeSigma. 0.1% DMSO (D8418, MilliporeSigma) served as the vehicle control in all experiments.

### Chemical screening

For the primary screen and secondary validation using the original ToxCast library stocks, three chorion-intact embryos at the sphere stage (3–4 hours post-fertilization, hpf) in a volume of 250 μL 1x E3 medium were transferred into each well of assay-ready 96-well plates containing pre-dispensed ToxCast chemicals, resulting in a final chemical concentration of 2 μM. Embryos were maintained at 28.5°C and manually evaluated for developmental toxicity at 1 and 2 days post-fertilization (dpf) on a dissecting microscope (Nikon SMZ745, Nikon SMZ25, or Zeiss Stemi 508).

Candidate developmental toxicants identified during the primary screen were independently re-tested using the original ToxCast library stocks under identical exposure conditions. Commercial concentration-response validation experiments were performed using embryos exposed beginning at shield stage (4 hpf) or 50% epiboly stage (5.25 hpf). Each treatment group consisted of 19–21 embryos, with three independent biological replicates derived from separate embryo clutches. Rescue experiments were performed under the same exposure paradigm beginning at the 50% epiboly stage, with 50 embryos per biological replicate (separate clutch).

Phenotypic endpoints were evaluated manually at both 1 and 2 days postfertilization (dpf) using a dissecting microscope. The endpoint scoring system was adapted from previous zebrafish developmental toxicity screens to maintain consistency with earlier ToxCast studies while expanding the phenotypic vocabulary to capture additional developmental abnormalities observed in the present screen (Truong et al. 2014).

### Calcium and magnesium rescue experiments

Rescue experiments were performed by supplementing 1× E3 medium with additional calcium chloride (CaCl₂) or magnesium sulfate (MgSO₄). Standard 1× E3 medium contains 0.33 mM Ca²⁺ and 0.33 mM Mg²⁺. Rescue media were prepared by supplementing E3 with an additional 50×, 70×, or 100× the basal calcium or magnesium concentration. The supplemented media were then used to prepare treatment solutions containing either 0.1% DMSO (vehicle control) or 3 μM purpurin.

Embryos at 50% epiboly (5.25 hours post-fertilization) were transferred to the treatment solutions and maintained at 28.5°C until 2 days post-fertilization (dpf). Embryos remained within the chorion during chemical exposure and were enzymatically dechorionated at 1 dpf using pronase to facilitate imaging and phenotypic assessment. At 2 dpf, representative images were acquired using a Nikon SMZ25 stereomicroscope equipped with a Hamamatsu ORCA-Flash4.0 digital CMOS camera, and developmental toxicity was assayed by morphological scoring using the same criteria described for the concentration-response validation experiments. Each treatment consisted of three independent biological replicates of 50 embryos.

### Imaging

Embryos from the primary high-throughput screen of 4,657 chemicals and the secondary validation of candidate developmental toxicants using the original ToxCast library stocks were imaged using a Nikon SMZ745 stereomicroscope. Representative images from the concentration-response validation experiments using commercially sourced chemicals (Fig. 3) were acquired with a Nikon SMZ745T stereomicroscope equipped with a Nikon DS-Fi3 camera. Rescue experiments were imaged using a Nikon SMZ25 stereomicroscope equipped with a Hamamatsu ORCA-Flash4.0 digital CMOS camera. Images were processed uniformly using Nikon NIS-Elements software, with identical acquisition settings applied to all embryos within each experiment.

### RNA extraction and RNAseq sample preparation

At 2 days post-fertilization (dpf), 20 live embryos exhibiting the characteristic treatment-induced developmental phenotype were collected for each biological replicate and transferred to RNase-free microcentrifuge tubes. (Thermo-Scientific Snap Cap Low Retention Microcentrifuge Tubes, 3434). Embryos were euthanized by rapid cooling on ice, after which residual treatment medium was carefully removed using 27 G×1.25 inch BD PrecisionGlide Needle (305109). Embryos were homogenized in 200 μL TRIzol reagent (ThermFisher-Scientific, 15596026) using a motorized pestle. An additional 800 μL TRIzol was added to each sample, followed by vortexing and incubation at room temperature for 5 min. Samples were centrifuged at 12,000 × *g* for 10 min at 4°C, and the clarified lysate was transferred to fresh tubes and stored at −80°C overnight.

Total RNA was extracted using the Direct-zol RNA Miniprep Kit (Zymo Research, 11-331) according to the manufacturer’s instructions. RNA concentration and purity were assessed using a NanoDrop 1000 spectrophotometer (Thermo Fisher Scientific), and RNA integrity was evaluated using an Agilent 4200 TapeStation System. Only samples with an A260/A280 ratio ≥2.0 and an RNA integrity number (RIN) ≥9.0 were used for library preparation. Directional poly(A)-selected mRNA libraries were prepared and sequenced by Novogene (Sacramento, CA, USA) using the NovaSeq X Plus platform to generate paired-end 150-bp reads.

### RNA sequencing analysis

RNA sequencing data were analyzed using the high-performance computing (HPC) cluster managed by the Biostatistics and Informatics Shared Resource (BISR) at Baylor College of Medicine. Raw FASTQ files were quality filtered and adapter trimmed using Trim Galore (v0.6.10) (Krueger, F. 2015). Trimmed reads were aligned to the zebrafish reference genome (GRCz11) using STAR (v2.7.1a) (Dobin et al. 2013), and gene-level read counts were generated with featureCounts (Liao et al. 2014). Count data were normalized using upper-quartile normalization in combination with the Remove Unwanted Variation (RUV) method (Risso et al. 2014). Differential gene expression analysis was performed using the edgeR R package (Robinson et al. 2010), and genes with a false discovery rate (FDR) <0.05 were considered significantly differentially expressed.

Gene set enrichment analysis (GSEA) was performed using GSEA v3.0 (https://www.gsea-msigdb.org/gsea/downloads.jsp) (Mootha et al. 2003; Subramanian et al. 2005). Ranked gene lists generated from the differential expression analysis were interrogated against the Gene Ontology Biological Process (GOBP), Kyoto Encyclopedia of Genes and Genomes (KEGG), Hallmark, Reactome, WikiPathways, and other Molecular Signatures Database (MSigDB) gene set collections. Pathways with an FDR *q*-value <0.25 were considered significantly enriched.

### Statistical Analysis

All statistical analyses and graphical representations were performed using GraphPad Prism (version 10.0 or later; GraphPad Software) and R Studio Version 2022.12.0+353. Concentration-response curves and EC₅₀ values were generated by nonlinear regression using log-transformed concentration data. Statistical comparisons for the concentration-response validation and rescue experiments were performed using one-way analysis of variance (ANOVA) followed by Dunnett’s multiple comparisons test, with each treatment group compared with the corresponding vehicle control. Data are presented as the mean ± standard deviation (SD) of three independent biological replicates unless otherwise indicated. Adjusted *P* values <0.05 were considered statistically significant.

## DATA AVAILABILITY

All data are contained within the paper and Supplementary Materials. The raw RNA-sequencing FASTQ files and processed gene-expression data for the purpurin and 4-aminoazobenzene experiments have been deposited in the NCBI Gene Expression Omnibus (GEO) under accession number GSE 345664.

## ACKNOWLEDGEMENTS

We thank Kelli Kessler and the staff of the Baylor College of Medicine Zebrafish Research Facility for excellent animal care and husbandry. We thank Kaley Neugebauer and Ahmed Mohamed for assistance with the experiments. We also thank the Baylor College of Medicine Biostatistics and Informatics Shared Resource for computational support, and Clifford Stephan, Ph.D., Institute of Biosciences and Technology, Texas A&M University, Houston, Texas, for assistance with automated liquid dispensing. Automated liquid dispensing was performed with support from the Institute of Biosciences and Technology Core Facility, supported by the Cancer Prevention and Research Institute of Texas (CPRIT) under award RP200668 (Research Resource Identifier: RRID). The Baylor College of Medicine high-performance computing cluster, managed by the Biostatistics and Informatics Shared Resource, is supported by grants from NCI P30-CA125123 and by institutional funds from the Dan L Duncan Comprehensive Cancer Center and Baylor College of Medicine. Figures were created in part with BioRender.com. ChatGPT (OpenAI GPT-5.5) was used to assist with R code generation and manuscript language editing. All scientific analyses, interpretation of results, and final manuscript content were reviewed and approved by the authors.

---

**Supplementary Figure 1:**
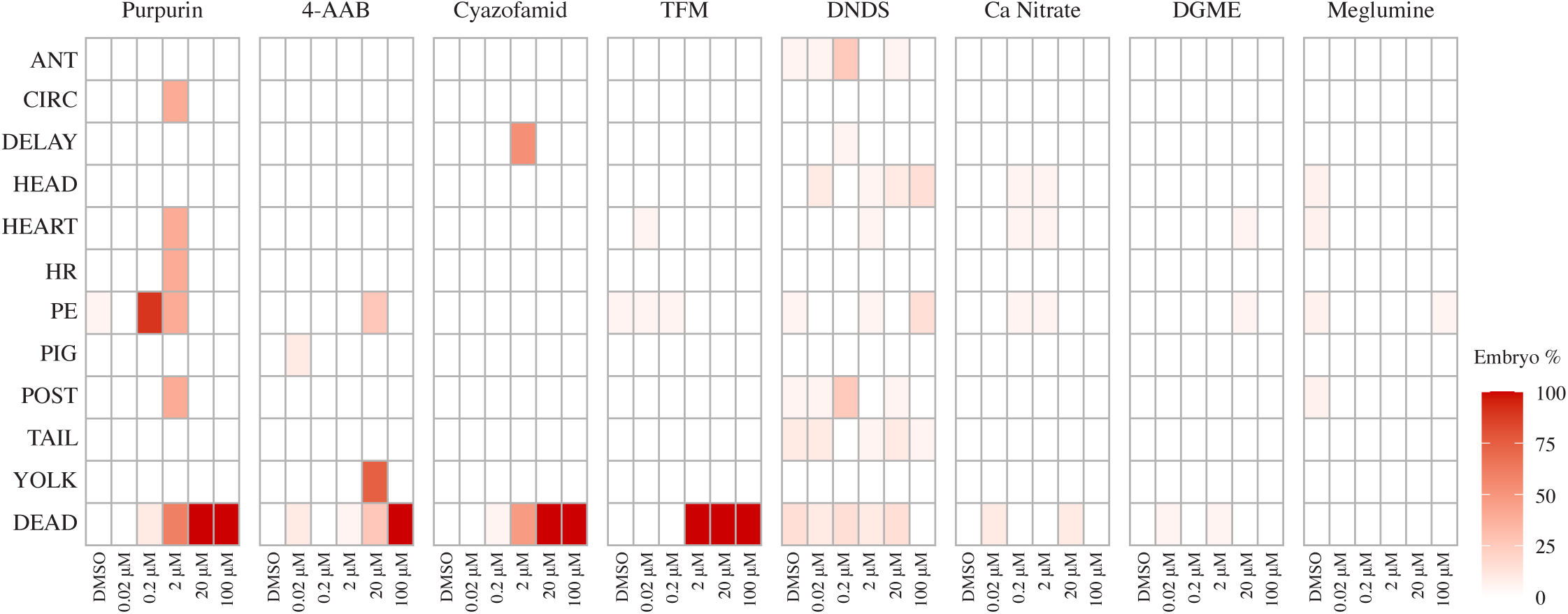
Phenotypic profiles of developmental abnormalities observed during concentration-response validation using commercially sourced chemicals. Heatmaps show the percentage of embryos exhibiting individual phenotypic endpoints at 2 dpf for each compound and concentrations tested in Figure 2B. Color intensity corresponds to the percentage of embryos displaying a given phenotype within each treatment group. Phenotypic endpoints include anterior defects (ANT), blood circulation defects (CIRC), developmental delay (DELAY), head abnormalities (HEAD), heart morphology defects (HEART), abnormal heart rate (HR), pericardial edema (PE), pigmentation defects (PIG), posterior truncation or thickening (POST), kinked or curly tail (TAIL), discolored yolk (YOLK), and mortality (DEAD). These data provide a phenotype-level breakdown of the abnormal embryo category summarized in Figure 2B.

**Supplementary Figure 2.**
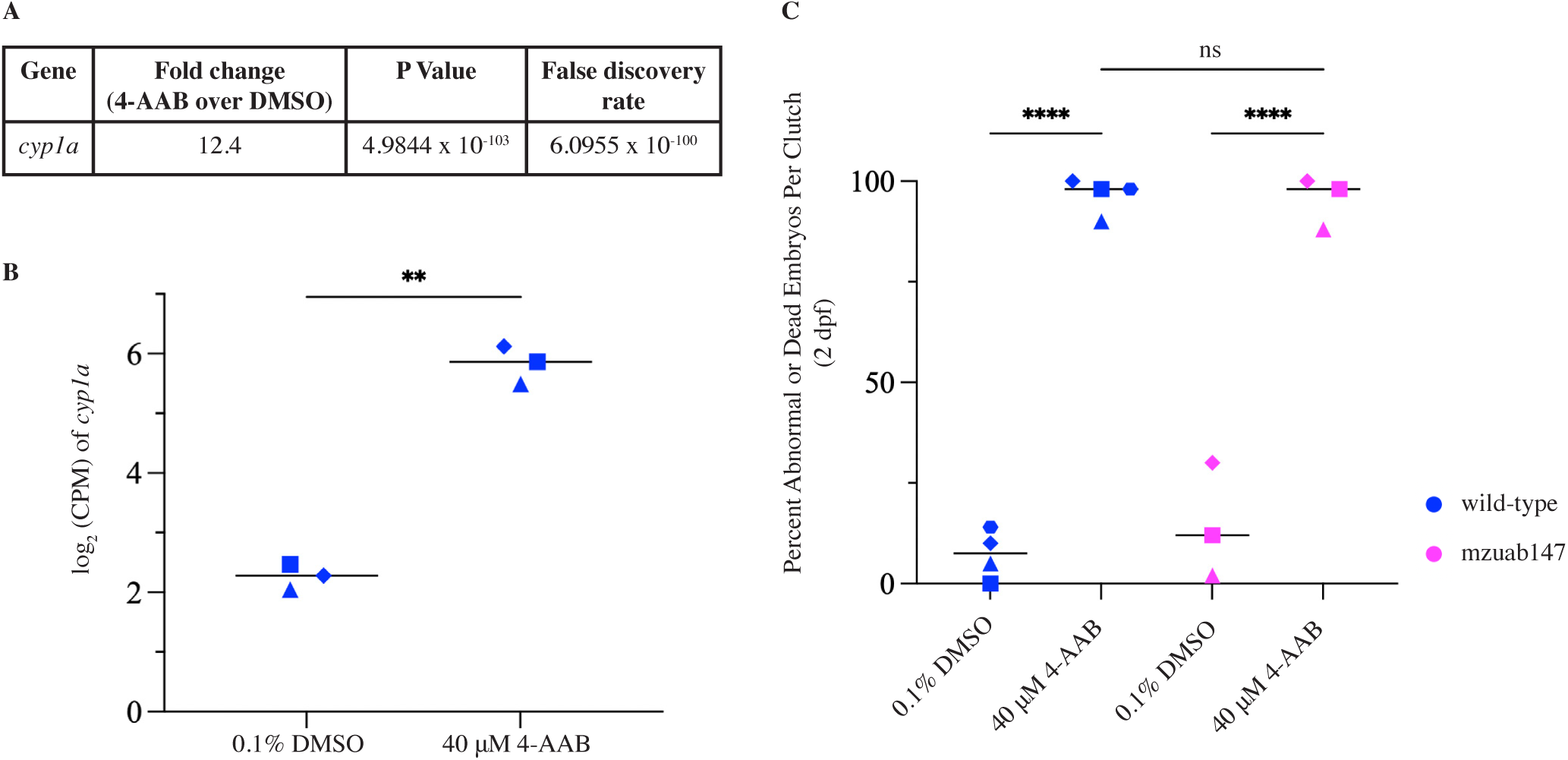
AHR2 is not required for 4-aminoazobenzene-induced developmental toxicity. **(A)** Differential expression of *cyp1a* in 4-aminoazobenzene (4-AAB)-treated zebrafish embryos compared with DMSO-treated controls, showing fold change, P value, and false discovery rate (FDR). **(B)** *cyp1a* expression, shown as log_2_ counts per million (CPM), in embryos treated with 0.1% DMSO or 40 µM 4-AAB. Points represent individual clutches (50 embryos per treatment group per clutch); horizontal lines indicate the mean. Groups were compared using a paired t-test. **(C)** Percentage of abnormal or dead wild-type and maternal-zygotic *ahr2* (mzuab147) mutant embryos following treatment with 0.1% DMSO or 40 µM 4-AAB. Loss of AHR2 function did not reduce 4-AAB-induced developmental toxicity. Each point represents an independent clutch (10–50 embryos per treatment group per clutch), and horizontal lines indicate the mean. Statistical comparisons were performed using a mixed-effects model. (**P < 0.01; ****P < 0.0001; ns, not significant)

**Supplementary Figure 3.**
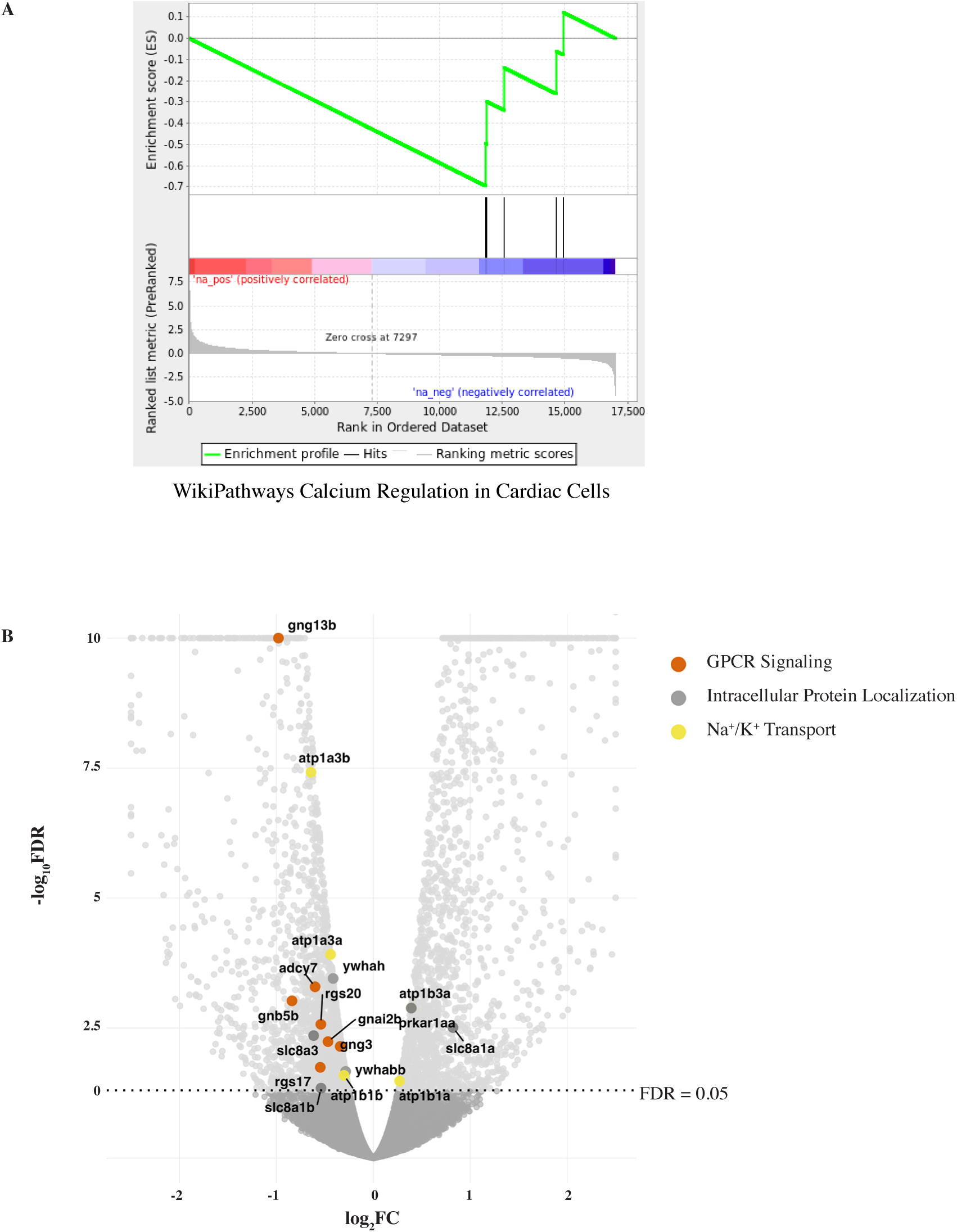
Enrichment and differential expression of genes associated with the WikiPathways Calcium Regulation in Cardiac Cells pathway following purpurin exposure. **(A)** Gene set enrichment analysis (GSEA) plot using a ranked transcriptomic dataset from purpurin-treated zebrafish embryos. The enrichment score (ES) is shown across the ranked gene list, with vertical black lines indicating the positions of pathway-associated genes. **(B)** Volcano plot showing differential gene expression following purpurin exposure. Each point represents a gene plotted by log fold change (FC) and −log (FDR). Genes associated with the functional groups: GPCR signaling, intracellular protein localization, and Na^+^/K^+^ transport are highlighted. The horizontal dotted line indicates the significance threshold (FDR = 0.05).

## References

Balik-Meisner M, Truong L, Scholl EH, La Du JK, Tanguay RL, Reif DM. 2018. Elucidating Gene-by-Environment Interactions Associated with Differential Susceptibility to Chemical Exposure. Environ Health Perspect 126:067010; doi:10.1289/EHP2662.

Birceanu O, McClelland GB, Wang YS, Brown JCL, Wilkie MP. 2011. The lampricide 3-trifluoromethyl-4-nitrophenol (TFM) uncouples mitochondrial oxidative phosphorylation in both sea lamprey (Petromyzon marinus) and TFM-tolerant rainbow trout (Oncorhynchus mykiss). Comp Biochem Physiol C Toxicol Pharmacol 153:342–349; doi:10.1016/j.cbpc.2010.12.005.

Clementi C, Nowik W, Romani A, Cibin F, Favaro G. 2007. A spectrometric and chromatographic approach to the study of ageing of madder (Rubia tinctorum L.) dyestuff on wool. Analytica Chimica Acta 596:46–54; doi:10.1016/j.aca.2007.05.036.

Delclos KB, Tarpley WG, Miller EC, Miller JA. 1984. 4-aminoazobenzene and N,N-dimethyl-4-aminoazobenzene as equipotent hepatic carcinogens in male C57BL/6 X C3H/He F1 mice and characterization of N-(Deoxyguanosin-8-yl)-4-aminoazobenzene as the major persistent hepatic DNA-bound dye in these mice. Cancer Res 44: 2540–2550.

Dix DJ, Houck KA, Martin MT, Richard AM, Setzer RW, Kavlock RJ. 2007. The ToxCast program for prioritizing toxicity testing of environmental chemicals. Toxicol Sci 95:5– 12; doi:10.1093/toxsci/kfl103.

Dobin A, Davis CA, Schlesinger F, Drenkow J, Zaleski C, Jha S, et al. 2013. STAR: ultrafast universal RNA-seq aligner. Bioinformatics 29:15–21; doi:10.1093/bioinformatics/bts635.

Gao M, Yang J, Wang Z, Yang B, Kuang H, Liu L, et al. 2016. Simultaneous Determination of Purpurin, Munjistin and Mollugin in Rat Plasma by Ultra High Performance Liquid Chromatography-Tandem Mass Spectrometry: Application to a Pharmacokinetic Study after Oral Administration of Rubia cordifolia L. Extract. Molecules 21:717; doi:10.3390/molecules21060717.

Green AJ, Mohlenkamp MJ, Das J, Chaudhari M, Truong L, Tanguay RL, et al. 2021. Leveraging high-throughput screening data, deep neural networks, and conditional generative adversarial networks to advance predictive toxicology. PLoS Comput Biol 17:e1009135; doi:10.1371/journal.pcbi.1009135.

Huerta B, Chung-Davidson Y-W, Bussy U, Zhang Y, Bazil JN, Li W. 2020. Sea lamprey cardiac mitochondrial bioenergetics after exposure to TFM and its metabolites. Aquat Toxicol 219:105380; doi:10.1016/j.aquatox.2019.105380.

Jäger I, Hafner C, Welsch C, Schneider K, Iznaguen H, Westendorf J. 2006. The mutagenic potential of madder root in dyeing processes in the textile industry. Mutation Research/Genetic Toxicology and Environmental Mutagenesis 605:22–29; doi:10.1016/j.mrgentox.2006.01.007.

Judson R, Richard A, Dix DJ, Houck K, Martin M, Kavlock R, et al. 2009. The toxicity data landscape for environmental chemicals. Environ Health Perspect 117:685–695; doi:10.1289/ehp.0800168.

Judson RS, Houck KA, Kavlock RJ, Knudsen TB, Martin MT, Mortensen HM, et al. 2010. In vitro screening of environmental chemicals for targeted testing prioritization: the ToxCast project. Environ Health Perspect 118:485–492; doi:10.1289/ehp.0901392.

Kojima M, Morita T, Degawa M, Hashimoto Y, Tada M. 1992. Differences in DNA damage induced by mutagenic and nonmutagenic 4-aminoazobenzene derivatives in Escherichia coli. Mutat Res 274:65–71; doi:10.1016/0921-8777(92)90044-4.

Krueger, F. 2015. Trim Galore!: A wrapper around Cutadapt and FastQC to consistently apply adapter and quality trimming to FastQ files, with extra functionality for RRBS data.

Lee J-H, Kim Y-G, Yong Ryu S, Lee J. 2016. Calcium-chelating alizarin and other anthraquinones inhibit biofilm formation and the hemolytic activity of Staphylococcus aureus. Sci Rep 6:19267; doi:10.1038/srep19267.

Liao Y, Smyth GK, Shi W. 2014. featureCounts: an efficient general purpose program for assigning sequence reads to genomic features. Bioinformatics 30:923–930; doi:10.1093/bioinformatics/btt656.

Lin C-F, Chen H-Y, Alalaiwe A, Hsiao Y-T, Jhong C-L, Chang S-H, et al. 2025. Harnessing Quinone Derivatives from *Rubia cordifolia* for Topical Therapy: Unveiling Structure– Activity and Structure–Permeation Relationships to Suppress Psoriasiform Inflammation. ACS Pharmacol Transl Sci 8:2270–2289; doi:10.1021/acsptsci.5c00352.

Macallum AB. 1925. THE PURPURIN METHOD OF LOCALIZING CALCIUM. Science 62:511; doi:10.1126/science.62.1614.511.

McCollum CW, Conde-Vancells J, Hans C, Vazquez-Chantada M, Kleinstreuer N, Tal T, et al. 2017. Identification of vascular disruptor compounds by analysis in zebrafish embryos and mouse embryonic endothelial cells. Reprod Toxicol 70:60–69; doi:10.1016/j.reprotox.2016.11.005.

Mitani S, Araki S, Takii Y, Ohshima T, Matsuo N, Miyoshi H. 2001. The Biochemical Mode of Action of the Novel Selective Fungicide Cyazofamid: Specific Inhibition of Mitochondrial Complex III in Phythium spinosum. Pesticide Biochemistry and Physiology 71:107–115; doi:10.1006/pest.2001.2569.

Mootha VK, Lindgren CM, Eriksson K-F, Subramanian A, Sihag S, Lehar J, et al. 2003. PGC-1alpha-responsive genes involved in oxidative phosphorylation are coordinately downregulated in human diabetes. Nat Genet 34:267–273; doi:10.1038/ng1180.

National Center for Biotechnology Information (2026). PubChem Compound Summary for CID 6051, 4-Aminoazobenzene. Available: https://pubchem.ncbi.nlm.nih.gov/compound/4-Aminoazobenzene [accessed 25 June 2026].

Niblett PD, Ballantyne JS. 1976. Uncoupling of oxidative phosphorylation in rat liver mitochondria by the lamprey larvicide TFM (3-trifluoromethyl-4-nitrophenol). Pesticide Biochemistry and Physiology 6:363–366; doi:10.1016/0048-3575(76)90046-8.

Padilla S, Corum D, Padnos B, Hunter DL, Beam A, Houck KA, et al. 2012. Zebrafish developmental screening of the ToxCast^TM^ Phase I chemical library. Reproductive Toxicology 33:174–187; doi:10.1016/j.reprotox.2011.10.018.

Pardo-Martin C, Chang T-Y, Koo BK, Gilleland CL, Wasserman SC, Yanik MF. 2010. High-throughput in vivo vertebrate screening. Nat Methods 7:634–636; doi:10.1038/nmeth.1481.

Park H, Shim JS, Kim BS, Jung HJ, Huh T-L, Kwon HJ. 2014. Purpurin inhibits adipocyte-derived leucine aminopeptidase and angiogenesis in a zebrafish model. Biochem Biophys Res Commun 450:561–567; doi:10.1016/j.bbrc.2014.06.017.

Peterson RT, Link BA, Dowling JE, Schreiber SL. 2000. Small molecule developmental screens reveal the logic and timing of vertebrate development. Proc Natl Acad Sci U S A 97:12965–12969; doi:10.1073/pnas.97.24.12965.

Raftery TD, Isales GM, Yozzo KL, Volz DC. 2014. High-Content Screening Assay for Identification of Chemicals Impacting Spontaneous Activity in Zebrafish Embryos. Environ Sci Technol 48:804–810; doi:10.1021/es404322p.

Reif DM, Truong L, Mandrell D, Marvel S, Zhang G, Tanguay RL. 2016. High-throughput characterization of chemical-associated embryonic behavioral changes predicts teratogenic outcomes. Arch Toxicol 90:1459–1470; doi:10.1007/s00204-015-1554-1.

Risso D, Ngai J, Speed TP, Dudoit S. 2014. Normalization of RNA-seq data using factor analysis of control genes or samples. Nat Biotechnol 32:896–902; doi:10.1038/nbt.2931.

Robinson MD, McCarthy DJ, Smyth GK. 2010. edgeR: a Bioconductor package for differential expression analysis of digital gene expression data. Bioinformatics 26:139–140; doi:10.1093/bioinformatics/btp616.

Rosenbaum DM, Rasmussen SGF, Kobilka BK. 2009. The structure and function of G-protein-coupled receptors. Nature 459:356–363; doi:10.1038/nature08144.

Sasaki YF, Izumiyama F, Nishidate E, Matsusaka N, Tsuda S. 1997. Detection of rodent liver carcinogen genotoxicity by the alkaline single-cell gel electrophoresis (Comet) assay in multiple mouse organs (liver, lung, spleen, kidney, and bone marrow). Mutat Res 391:201–214; doi:10.1016/s1383-5718(97)00072-7.

Subramanian A, Tamayo P, Mootha VK, Mukherjee S, Ebert BL, Gillette MA, et al. 2005. Gene set enrichment analysis: a knowledge-based approach for interpreting genome-wide expression profiles. Proc Natl Acad Sci U S A 102:15545–15550; doi:10.1073/pnas.0506580102.

Thomas DG, Shankaran H, Truong L, Tanguay RL, Waters KM. 2019. Time-dependent behavioral data from zebrafish reveals novel signatures of chemical toxicity using point of departure analysis. Comput Toxicol 9:50–60; doi:10.1016/j.comtox.2018.11.001.

Truong L, Bugel SM, Chlebowski A, Usenko CY, Simonich MT, Simonich SLM, et al. 2016. Optimizing multi-dimensional high throughput screening using zebrafish. Reprod Toxicol 65:139–147; doi:10.1016/j.reprotox.2016.05.015.

Truong L, Reif DM, St Mary L, Geier MC, Truong HD, Tanguay RL. 2014. Multidimensional in vivo hazard assessment using zebrafish. Toxicol Sci 137:212–233; doi:10.1093/toxsci/kft235.

Westerfield, M. 2020. The Zebrafish Book. A Guide for the Laboratory Use of Zebrafish (Danio rerio). 4th ed. University of Oregon Press:Eugene, OR.

Yao Y, Sun S, Fei F, Wang J, Wang Y, Zhang R, et al. 2017. Screening in larval zebrafish reveals tissue-specific distribution of fifteen fluorescent compounds. Dis Model Mech 10:1155–1164; doi:10.1242/dmm.028811.

Zhang G, Marvel S, Truong L, Tanguay RL, Reif DM. 2016. Aggregate entropy scoring for quantifying activity across endpoints with irregular correlation structure. Reprod Toxicol 62:92–99; doi:10.1016/j.reprotox.2016.04.012.

Zhang G, Roell KR, Truong L, Tanguay RL, Reif DM. 2017a. A data-driven weighting scheme for multivariate phenotypic endpoints recapitulates zebrafish developmental cascades. Toxicol Appl Pharmacol 314:109–117; doi:10.1016/j.taap.2016.11.010.

Zhang G, Truong L, Tanguay RL, Reif DM. 2017b. A New Statistical Approach to Characterize Chemical-Elicited Behavioral Effects in High-Throughput Studies Using Zebrafish. PLoS One 12:e0169408; doi:10.1371/journal.pone.0169408.

